# Multi-enhancer transcriptional hubs confer phenotypic robustness

**DOI:** 10.1101/575175

**Authors:** Albert Tsai, Mariana RP Alves, Justin Crocker

## Abstract

We had previously shown in *Drosophila melanogaster* embryos that low-affinity Ultrabithorax (Ubx)-responsive *shavenbaby* (*svb*) enhancers drive robust expression using localized transcriptional environments and that active *svb* enhancers tended to colocalize, even when placed on different chromosomes (Tsai et al., 2017). Here, we test the hypothesis that these multi-enhancer “hubs” improve robustness by increasing transcription factor retention near transcription sites. Deleting a redundant enhancer from the *svb* locus led to reduced trichome numbers in embryos raised at elevated temperatures. Using high-resolution fluorescence microscopy, we observed lower Ubx concentration and transcriptional output in this deletion allele. Transcription sites of the full *svb cis*-regulatory region inserted into a different chromosome colocalized with the *svb* locus, increasing Ubx concentration, the transcriptional output of *svb*, and partially rescuing the phenotype. Thus, multiple enhancers could reinforce a local transcriptional hub to buffer against environmental stresses and genetic perturbations, providing a mechanism for phenotypical robustness.

**Impact statement:** Multiple enhancers in physical proximity can reinforce shared transcriptional “hubs” to retain transcription factors, providing a buffer during environmental stresses and genetic perturbations to preserve phenotypic robustness.

## Introduction

> “Come together, right now, over me”
>
> -The Beatles (who may, or may not, have been singing about transcription)

During embryogenesis, transcriptional regulation controls the expression of genes that determine the fate of specific cells, ultimately leading to the correct patterning of the animal body plan (Long et al., 2016; Mallo and Alonso, 2013; Reiter et al., 2017; Spitz and Furlong, 2012). This involves coordinating a complex series of interactions between transcription factors and their target binding sites on DNA, leading to the recruitment or exclusion of active RNA polymerases, which determines the transcriptional state of the gene. Live imaging experiments have shown that transcription factor binding in eukaryotic cell lines and embryos is dynamic but transient, occurring frequently but with each event lasting for at most a few seconds (Chen et al., 2014; Izeddin et al., 2014; Liu et al., 2014; Normanno et al., 2015). Additionally, recent studies have shown that many developmental enhancers harbor kinetically weak, low-affinity binding sites, required to achieve sufficient specificity amongst closely related transcription factors (Antosova et al., 2016; Crocker et al., 2010, 2015a, 2015b, Farley et al., 2015, 2016; Gaudet and Mango, 2002; Lebrecht et al., 2005; Lorberbaum et al., 2016; Rister et al., 2015; Rowan et al., 2010; Tanay, 2006). One example is the Homeobox (Hox) family that is responsible for body segment identity along the anterior-posterior axis in animals. Because Hox transcription factors descent from a common ancestor, their preferences for binding sequences are very similar (Berger et al., 2008; McGinnis and Krumlauf, 1992; Noyes et al., 2008). To achieve specificity, several enhancers in the *shavenbaby* (*svb*) locus make exclusive use of low-affinity binding sequences for the Hox factor Ultrabithorax (Ubx) (Crocker et al., 2015a). Svb is a transcription factor that drives the formation of trichomes, epidermal projections on the surface of the segmented fly embryo (Chanut-Delalande et al., 2006; Delon et al., 2003; Payre et al., 1999). Thus, a key question was how kinetically inefficient binding sequences are able to achieve robust transcriptional activation in developing embryos.

We have previously shown in living *Drosophila melanogaster* embryos that Ubx transiently but repeatedly explores the same physical locations in a nucleus, which are likely hotspots with clusters of binding sites (Tsai et al., 2017). We have additionally shown that transcriptional microenvironments of high local Ubx and cofactor concentrations surround active transcription sites driven by low-affinity *svb* enhancers. As the distributions of many transcription factors in the nucleus are highly heterogeneous, the transcriptional activity of low-affinity enhancers depends on the local microenvironments. Interestingly, we observed that transcriptionally active, minimalized versions of two of the three ventral *svb* enhancers (*E3* and *7*) are Ubx-responsive and preferentially appear near or overlap spatially with transcription sites of the endogenous *svb* gene, despite being on different chromosomes (Crocker et al., 2015a; Tsai et al., 2017). This colocalization suggests microenvironments could be shared between related enhancers to increase transcriptional output, where enhancers would synergistically form a larger local trap for transcription factors than each could alone. Retaining multiple enhancers within a microenvironment would also provide redundancy in case individual enhancers are compromised and buffer negative impacts when the system is subjected to stress. This idea is consistent with the observed phenotypical resilience of *svb* enhancers under temperature stress (Crocker et al., 2015a; Frankel et al., 2010). These results would also be consistent with multi-component transcriptional “hubs” (Boija et al., 2018; Cisse et al., 2013; Furlong and Levine, 2018; Ghavi-Helm et al., 2014; Lim et al., 2018; Mir et al., 2017, 2018).

To understand the mechanistic implications of having multiple enhancers in a shared microenvironment, here we examined the transcriptional robustness of the *svb* locus to temperature-induced stress in flies harboring the wild-type allele or one containing a deletion (*Df(X)svb*^*108*^) that includes the ventral *svb* enhancer *DG3*. When embryos were raised at high temperatures, we observed phenotypical defects in ventral trichome formation for the *DG3*-deletion *svb* allele but not for the wild-type. At the molecular level, Ubx concentrations around transcription sites of the *DG3*-deletion allele decreased. The transcriptional output of *svb* without *DG3* also decreased. To test the hypothesis that shared microenvironments modulate transcriptional output and provide buffering under stress, we sought to rescue the *DG3*-deletion allele through inserting the complete *svb* cis-regulatory region on a BAC (*svbBAC*) on a different chromosome. We observed that Ubx concentration around active transcription sites of the *DG3*-deletion allele and their transcriptional output increased when the *svbBAC* is physically nearby. Moreover, we found that trichome formation was partially rescued at high temperature. As a result, our findings support the hypothesis that shared microenvironments provide a mechanism for phenotypic robustness.

## Results

### The *DG3* enhancer responds specifically to Ubx in the A1 segment

The ventral *svb* enhancers *DG3, E3* and *7* (Figure 1A) contain low-affinity Ubx binding sites and have been shown to be transcribed in microenvironments of high Ubx concentrations in the first abdominal (A1) segment on the ventral surface of the embryo (Tsai et al., 2017). Each of these enhancers produces ventral stripes of expression along segments A1-A7 in the embryo, resembling the endogenous expression pattern of *svb* (Figure 1B). Each enhancer contributes to different but partially overlapping portions of the total expression pattern. Furthermore, they have different Ubx ChIP enrichment profiles (Figure 1-figure supplement 1). Whereas the interaction of *E3* and *7* with Ubx had been previously explored in detail (Crocker et al., 2015a), *DG3* remained unexplored. Therefore, we tested the response of the *DG3* enhancer to Ubx by altering Ubx levels and measuring the transcriptional output with a reporter gene (*lacZ*). In wild-type embryos, the *DG3* reporter gene was expressed ventrally in stripes along segments A1-A7, in addition to narrow thoracic stripes (Figure 1C). In the absence of Ubx, *DG3* reporter expression was almost completely lost on the ventral side of A1 and significantly reduced between A2-A7 (Figure 1D), consistent with the responses of *E3, 7* and the full *svb* locus (Crocker et al., 2015a). Ubiquitous expression of Ubx increased the expression levels in A1-A7, in addition to generating ectopic expressions in the thoracic segments T1-T3 and A8 (Figure 1E). In summary, we showed that *DG3* responds specifically to Ubx for driving gene expression, which is consistent with our previous observation of the localization of *DG3*-driven transcription sites within Ubx microenvironments (Tsai et al., 2017).

**Figure 1.**
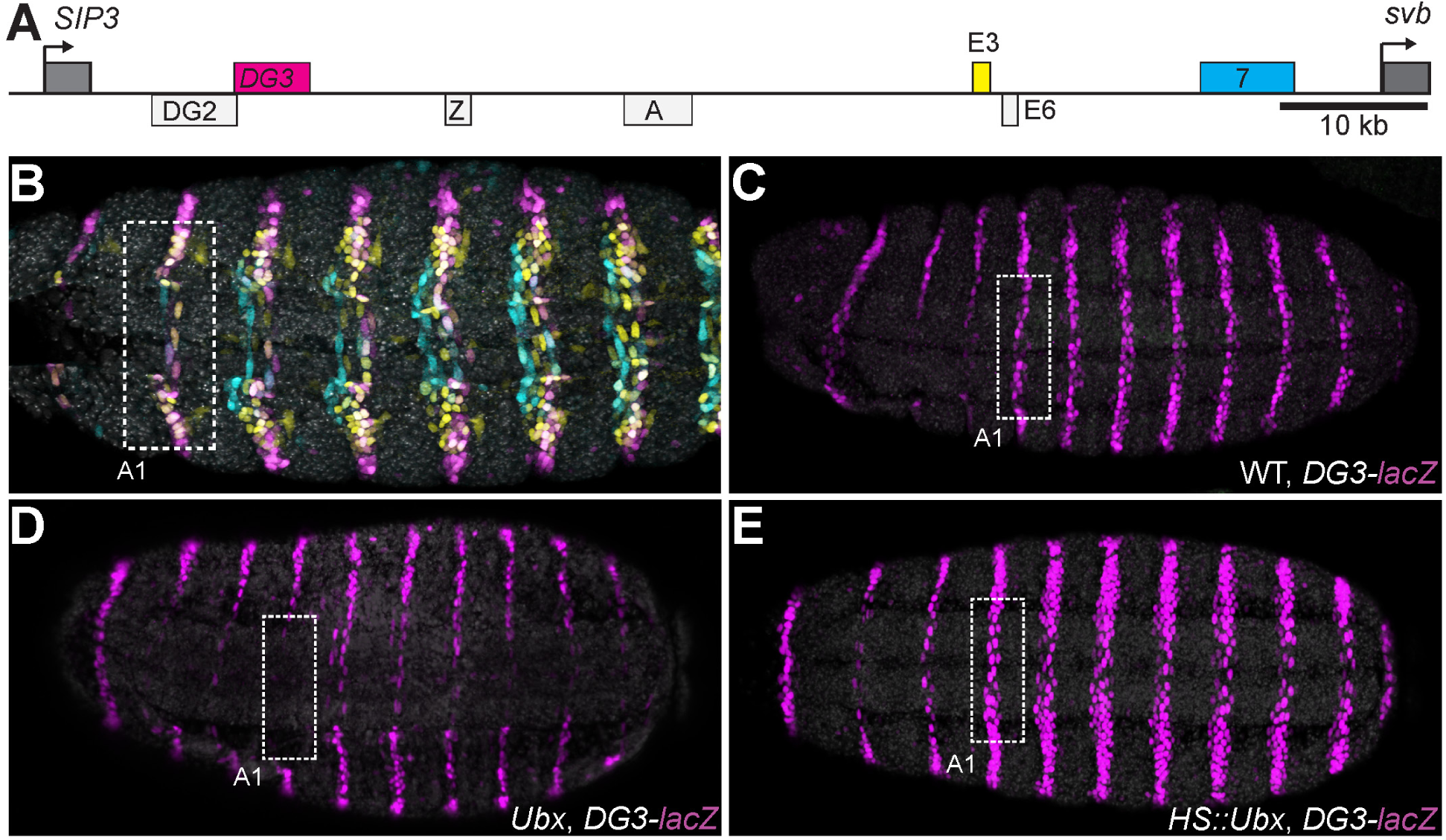
Ubx drives the expression of the *DG3 shavenbaby* enhancer along the ventral abdominal segments. (A) The *cis*-regulatory region of the *shavenbaby* (*svb*) gene contains 3 enhancers expressing stripes on the ventral side of the abdominal segments: *DG3, E3* and *7*. (B) The expression patterns of the 3 enhancers are partially overlapping. The color scheme corresponds to (A), where *DG3* is magenta, *E3* is yellow and *7* is cyan. (C) Expression pattern of a reporter construct with the *DG3* enhancer driving LacZ expression in an embryo with wild-type Ubx expression, as visualized using immunofluorescence staining. (D) *DG3* reporter in Ubx null mutant shows no expression in A1 and significantly weakened expression in the other abdominal segments. (E) Overexpression of Ubx driven through a heat shock promoter induces overexpression of *DG3* reporter in all abdominal segments.

### Deletion of a region including *DG3* enhancer causes defects in ventral trichome formation specifically at elevated temperatures

Given the strong ventral stripes that *DG3* generated in the abdominal segments, we next explored the phenotypical impact of its activity in driving trichome formation. It has been previously shown that deleting a region in the *svb* locus containing *DG3* (*Df(X)svb*^*108*^) leads to reduced phenotypic robustness of *svb* under non-optimal temperatures (Frankel et al., 2010). This *svb DG3*-deletion allele encompasses the enhancers *DG2, DG3* and *Z* (Figure 2A)—of which only *DG3* is a ventral enhancer.

**Figure 2.**
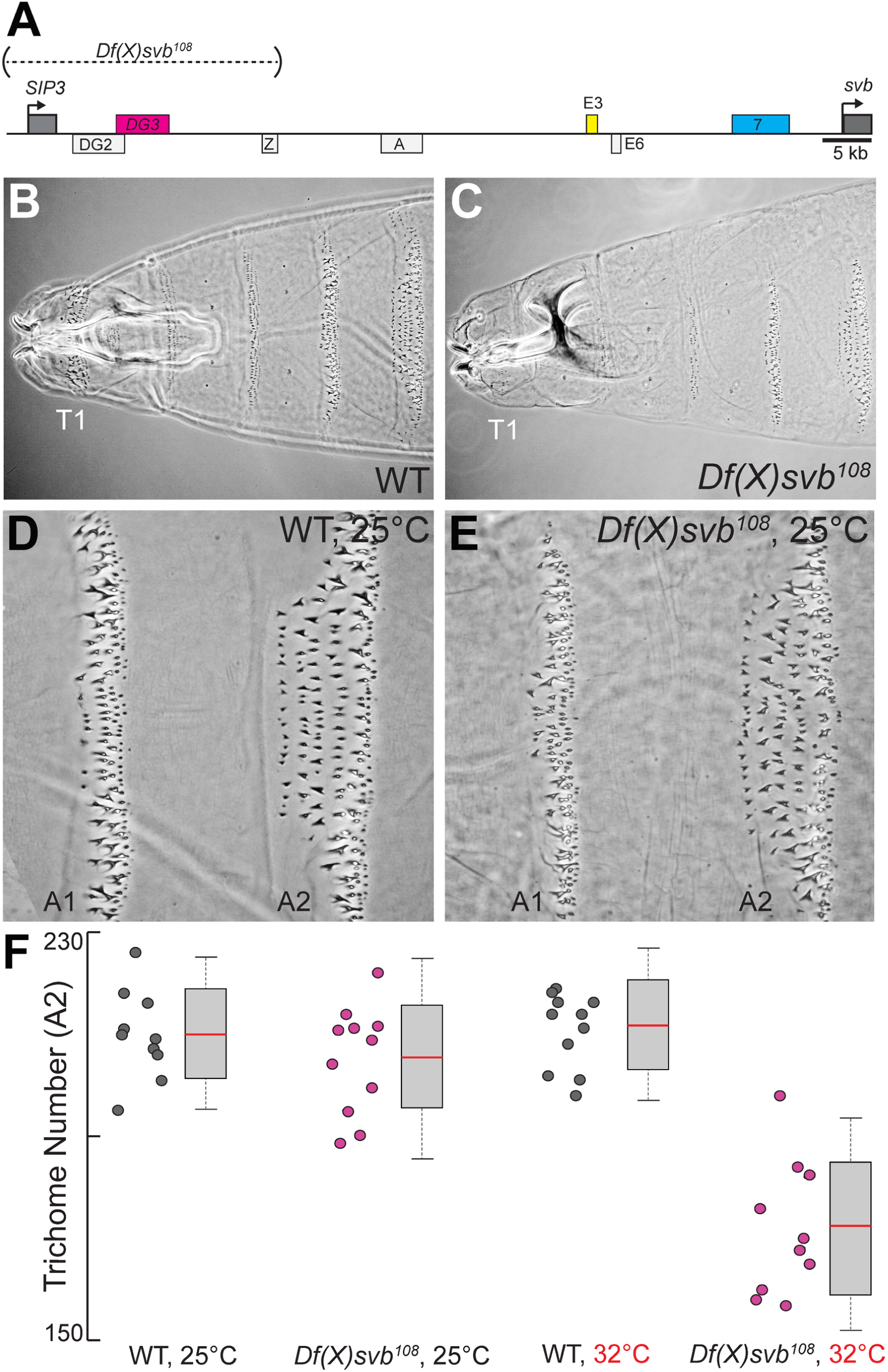
Deletion of a region from the *svb* locus containing *DG3* reduces ventral trichome numbers under heat-induced stress. (A) The *Df(X)svb*^*108*^ allele contains a deletion in the *cis*-regulatory of svb spanning 3 enhancers: *DG2, DG3* and *Z*. Of those, only *DG3* expresses on the ventral side. (B) A cuticle preparation of a wild-type (w^1118^) larva. (C) A cuticle preparation of a larva carrying *Df(X)svb*^*108*^. The lack of trichomes along the T1 segment is a recessive marker used in subsequent experiments to select for embryos/larvae carrying this deletion allele. (D) Wild-type phenotype of trichomes along the A1 and A2 segments at 25 °C. (E) At 25 °C, the *Df(X)svb*^*108*^ deletion allele did not show a clear mutant phenotype along the A1 and A2 segments. (F) Deficiencies of the deletion allele only become clear when the animal is subjected to elevated temperature at 32 °C, showing reduced trichome numbers in the A2 segment. The number of larvae counted was: 10 for wild-type at 25 °C, 11 for *Df(X)svb*^*108*^ at 25 °C, 11 for wild-type at 32 °C and 10 for *Df(X)svb*^*108*^ at 32 °C. In box plots, center line is mean, upper and lower limits are standard deviation and whiskers show 95% confidence intervals.

In the A1 and A2 segments at 25 °C, deletion of the *DG3* enhancer did not result in a clear change in ventral trichome formation in the abdominal segments (Figure 2B-E), perhaps due to the redundancy provided by overlapping expression patterns from other *svb* enhancers. However, the T1 trichomes were missing in larvae homozygous for the deletion (*Df(X)svb*^*108*^) allele (compare Figure 2B and 2C), which we subsequently used as a recessive marker to select for embryos carrying the deletion allele when crossing *Df(X)svb*^*108*^ flies to other lines. Also, we observed defects in trichome formation in the dorsal edges of the stripe pattern, which are exclusively covered by *DG3* (Figure 2-figure supplement 1A-C). This is consistent with a lack of redundancy in enhancer usage in these areas. The trichome number in regions covered by the overlapping expression of the *E3, 7* and *DG3* enhancers in segment A2 did not significantly reduce at 25 °C upon the deletion of *DG3* (Figure 2D & E). However, larvae homozygous for the *Df(X)svb*^*108*^ allele developed at 32 °C produced fewer trichomes compared to wild-type flies (Figure 2F). These results are similar to those shown with quartenary A5 trichomes (Frankel et al., 2010). However, the mechanisms behind this loss of phenotypic robustness under heat-induced stress are yet to be understood in detail.

### Transcription sites from the DG3-deletion allele have weaker Ubx microenvironment and lower transcriptional output

To address the mechanistic causes leading to the reduced number of ventral trichomes we observed for the *Df(X)svb*^*108*^ deletion allele, we imaged Ubx distributions and the transcriptional output of the *svb* gene in fixed *Drosophila melanogaster* embryos using high-resolution confocal microscopy. We reasoned that the defect could be with changes in the Ubx concentration around the enhancers (input) and/or the transcriptional output of the gene (output). The samples were stained with immunofluorescence (IF) for Ubx and RNA fluorescence *in situ* hybridization (FISH) for *svb* transcription sites as previously described (Tsai et al., 2017). We imaged both embryos containing the wild-type *svb* allele and the *Df(X)svb*^*108*^ allele, raised at either 25 °C or 32 °C.

To gauge the Ubx concentration around a transcription site, we counted the averaged intensity in the Ubx IF channel within a 40×40 pixel (2.8×2.8 µm) box centered on the transcription site (Figure 3A & B, see “Analysis of microenvironment and *svb* transcription intensity” in materials and methods). In nuclei from the A1 segment, Ubx distributions around *svb* transcription sites with the wild-type allele did not change between 25 °C and 32 °C (Figure 3C). Transcription sites in embryos with the *DG3*-deletion (*Df(X)svb*^*108*^) allele had a local Ubx concentration that is indistinguishable from wild-type at 25 °C (Figure 3B, left panel, and 3C). However, there was a moderate decrease in Ubx intensity compared to the wild-type when we subjected the *DG3*-deletion embryos to heat-stress (Figure 3B, right panel, and 3C). To measure the transcriptional output of *svb*, we adopted the same approach, but in the *svb* RNA FISH channel (Figure 3D). Interestingly, we detected clear decreases in transcriptional output when the embryos are heat-stressed at 32 °C, even with the wild-type allele. The *Df(X)svb*^*108*^ allele at 25 °C showed reduced levels of transcriptional output noticeably lower than the wild-type under heat-shock. At 32 °C, the transcriptional output further decreased in the mutant. In sum, stress conditions impact the transcriptional output of enhancers even if the Ubx input did not change considerably for both the wild-type and the deletion mutant.

**Figure 3.**
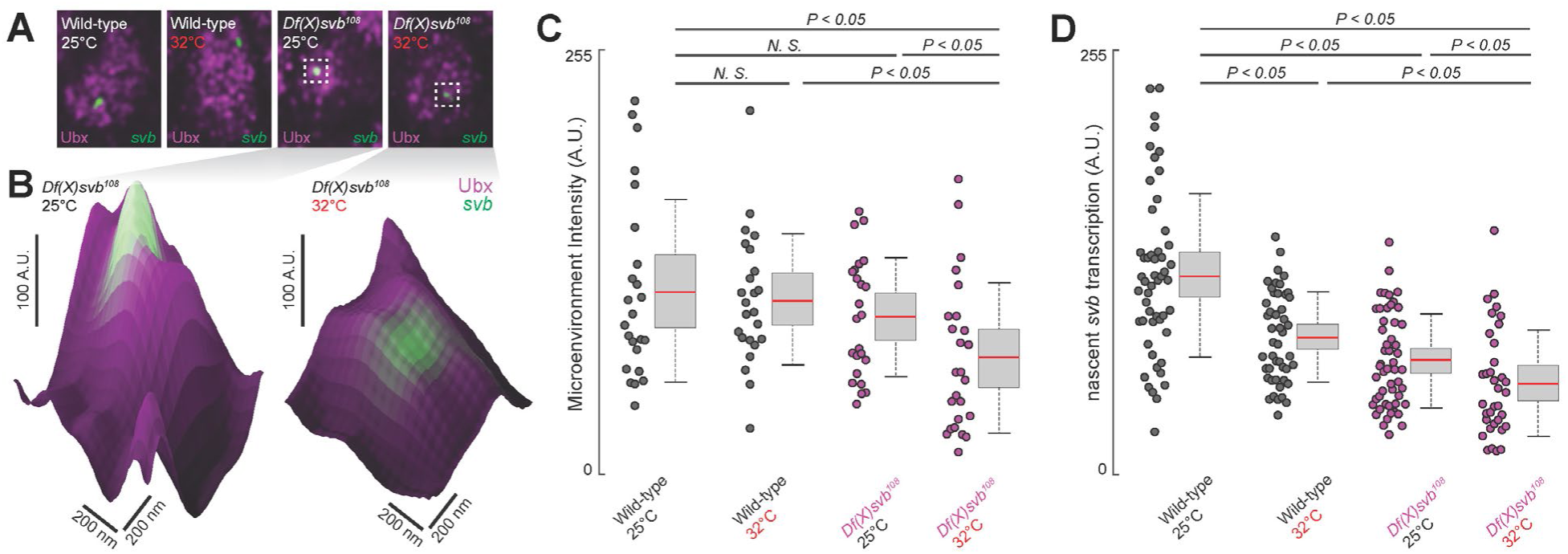
Deletion of the *cis*-region of *svb* containing *DG3* led to defects in the Ubx microenvironment and *svb* transcriptional output. (A) Panels showing a nucleus from embryos with either the wild-type (w^1118^) or *Df(X)svb*^*108*^ deletion allele at *svb* and either at normal (25 °C) or elevated temperature (32 °C), imaged using confocal fluorescence microscopy. Ubx (shown in magenta) is stained using immunofluorescence (IF) and the *svb* transcription sites (shown in green) are stained using fluorescence *in situ* hybridization (FISH). (B) Zoomed-in panes centered on *svb* transcription sites, with the height of the surface plots representing the Ubx intensity. (C) Integrating the Ubx intensity surrounding transcription sites shows a moderate defect in the Ubx concentration around the deletion allele, but only at elevated temperature. The number of nuclei quantified was: 24 for wild-type at 25 °C, 24 for wild-type at 32 °C, 24 for *Df(X)svb*^*108*^ at 25 °C and 25 for Df(X)svb^108^ at 32 °C. (D) The integrated intensity of *svb* transcriptional output shows that there is a drop in transcriptional output for the deletion allele compared to the wild-type at both 25 and 32 °C. Interestingly, even the wild-type showed reduced transcriptional output at elevated temperature (32 °C). The number of transcription sites quantified was: 52 for wild-type at 25 °C, 46 for wild-type at 32 °C, 51 for *Df(X)svb*^*108*^ at 25 °C and 35 for Df(X)svb^108^ at 32 °C. Note data sets in (C) and (D) were analyzed separately. We analyzed 3 embryos for each genotype/temperature combination. One-sided Student’s *t*-test was applied for each individual comparison. In box plots, center line is mean, upper and lower limits are standard deviation and whiskers show 95% confidence intervals.

### *Df(X)svb*^*108*^ deficiencies are rescued upon insertion of the full *svb cis*-regulatory region in a different chromosome

Having observed in the past that transcriptional microenvironments can be shared between related *svb* enhancers on different chromosomes (Tsai et al., 2017), we wondered whether this phenomenon could enhance transcriptional output and thus buffer against adverse environmental conditions. Therefore, we tested the capacity of a DNA sequence containing the full *svb cis*-regulatory region to rescue the described molecular and developmental defects of the *Df(X)svb*^*108*^ allele. For this purpose, we used a transgenic fly line, where a bacterial artificial chromosome (BAC) carrying the complete *cis*-regulatory region of *svb* (Preger-Ben Noon et al., 2018a) was integrated into chromosome 2. To exclude *svb* mRNA from effecting the rescue, this *svbBAC* construct drives a *dsRed* reporter gene instead of another copy of *svb*. We confirmed that DsRed protein expression driven by this regulatory sequence recapitulates the *svb* expression patterns (Figure 4A) in *D. melanogaster* embryos and is responsive to Ubx—the lack of Ubx leads to a decrease of expression in the A1 segment (Figure 4B).

**Figure 4.**
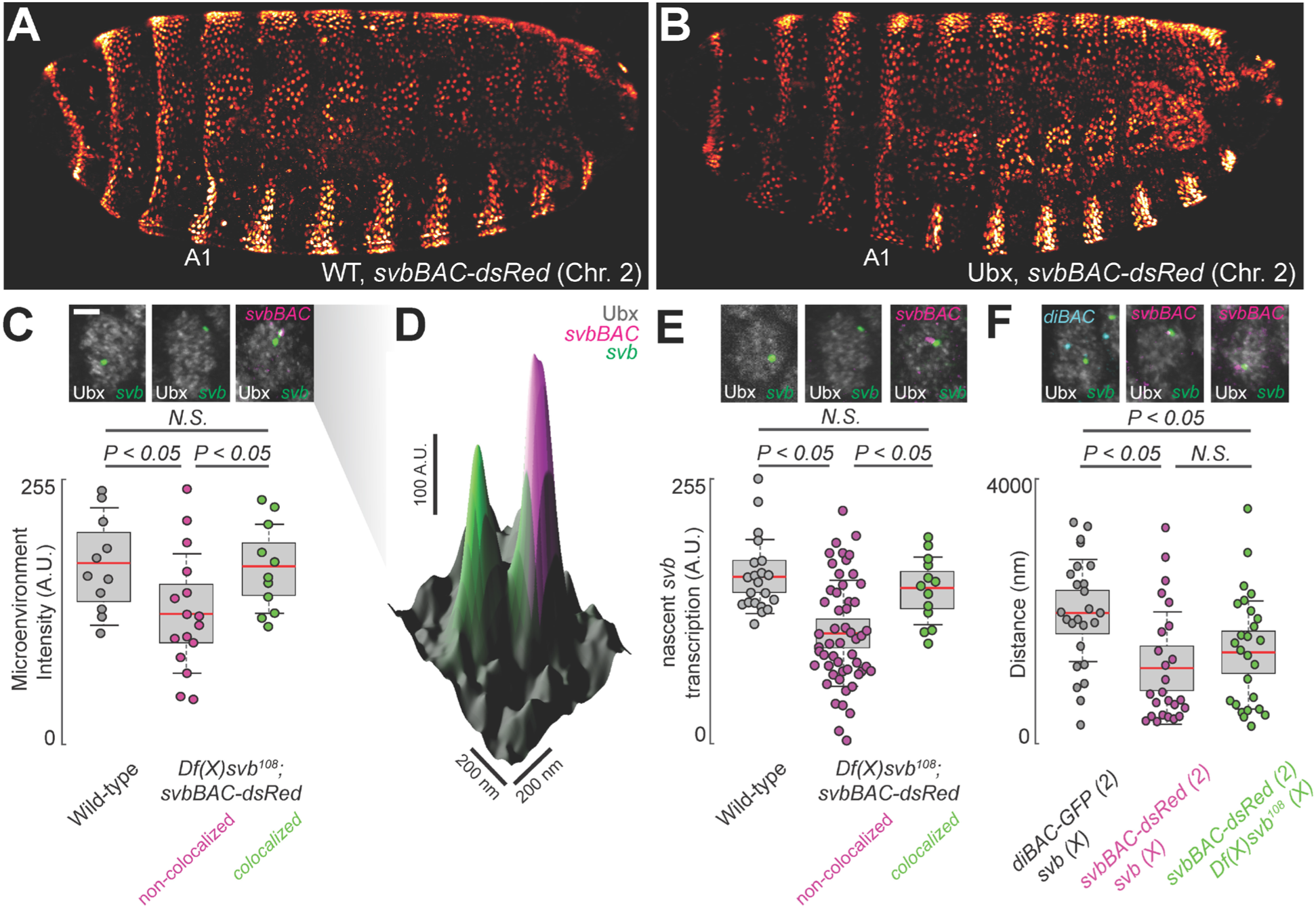
Introduction of the *cis*-regulatory region of *svb* on another chromosome rescues the microenvironment deficiencies of the *Df(X)svb*^*108*^ deletion mutant. (A & B) The *svbBAC* driving the expression of *dsRed* inserted into the second chromosome drives a similar expression pattern as the wild-type *svb* locus and responds similarly to Ubx. (C) At 32 °C, Ubx concentration around *svb* transcription sites recovered to wild-type levels in nuclei containing colocalized *svb* and *dsRed* transcription sites (colocalized) in *Df(X)svb*^*108*^ x *svbBAC-dsRed* embryos. Ubx levels around transcription sites of *svb* in the same embryos in nuclei without a *dsRed* transcription site (non-colocalized) did not recover. The number of *svb* transcription sties quantified was: 11 for wild-type, 16 for *Df(X)svb*^*108*^ not near a *dsRed* transcription site and 11 for *Df(X)svb*^*108*^ near a *dsRed* transcription site. (D) A surface plot (the height representing Ubx intensity) showing two *svb* transcription sites in a nucleus, with the one on the right overlapping with a *svbBAC-dsRed* transcription site and showing higher Ubx concentration. (E) The *svb* FISH intensity (representing transcriptional output) in *Df(X)svb*^*108*^ x *svbBAC-dsRed* embryos at 32 °C recovered to wild-type levels only when the *svb* transcription site is near a *dsRed* transcription site, similar to Ubx concentration. The number of transcription sites quantified was: 21 for wild-type, 53 for *Df(X)svb*^*108*^ not near a *dsRed* transcription site and 26 for *Df(X)svb*^*108*^ near a *dsRed* transcription site. (F) In nuclei having both *svb* (for both the wild-type and the *Df(X)svb*^*108*^ allele) and *svbBAC-dsRed* transcription sites, the distances between them is on average closer than between that of *svb* and a reporter construct of an unrelated gene, *diachete* (*diBAC-gfp*), inserted into the same location as *svbBAC* on the second chromosome. The pairs of distance quantified were: 25 between *diBAC-gfp* and wild-type *svb*, 25 between *svbBAC-dsRed* and wild-type *svb* and 26 between *svbBAC-dsRed* and *Df(X)svb*^*108*^. We analyzed 3 embryos for each genotype. One-sided Student’s *t*-test was applied for each individual comparison. In box plots, center line is mean, upper and lower limits are standard deviation and whiskers show 95% confidence intervals.

To test the rescue, *Df(X)svb*^*108*^ embryos or larvae with a *svbBAC-dsRed* crossed into them were incubated at 32 °C. We observed that the introduction of the *svb* regulatory region was able to rescue both molecular and functional defects observed from the loss of the region containing *DG3*. Both local Ubx concentration around *svb* transcription sites (Figure 4C & D) and their transcriptional output (Figure 4E) were restored to wild-type levels, but only when they co-localized with an active *svbBAC-dsRed* transcription site in the same nucleus. Wild-type (w^1118^) x *svbBAC-dsRed* embryos and larvae were identical to wild-type alone without any significant changes in trichome number and Ubx levels around *svb* transcription sites (Figure 4-figure supplement 1). We additionally observed that active *svbBAC* reporter gene transcription sites are close to *svb* transcription sites in embryos from crosses between *svbBAC-dsRed* and wild-type (w^1118^) flies (Figure 4F). This observation is also true for embryos from crosses between *svbBAC-dsRed* and *Df(X)svb*^*108*^ flies, suggesting that the co-localization of transcriptional microenvironments between related enhancers could occur even under stressed conditions. This effect was not observed for the unrelated regulatory region of *diachete* driving expression of GFP, which was inserted on a BAC in the same chromosomal location as *svbBAC* (Fig. 4F).

Regarding phenotype, ventral trichome formation on the A1 segment (Figure 5A-C), which is reduced with the *DG3*-deletion allele, is also rescued by the introduction of *svbBAC* (Figure 5D). Interestingly, the loss of the outer edge trichomes in A1 (in the black brackets, as shown in Figure 5A-C, where only *DG3* provides coverage) with the *DG3*-deletion allele was not rescued with *svbBAC*. Additionally, introducing only the *DG3* enhancer as opposed to *svbBAC* did not rescue trichome formation under heat-stress (Figure 5D).

**Figure 5.**
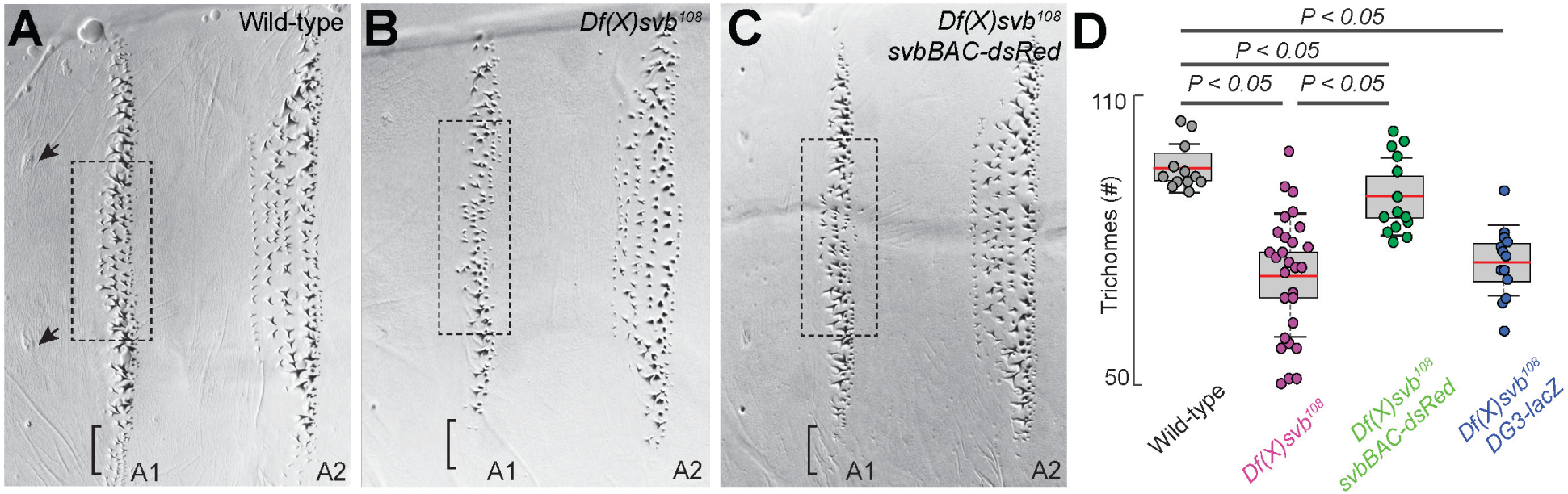
The complete *cis*-regulatory region of *svb* rescues trichome number. (A-C) Cuticle preparations from larvae with wild-type *svb, Df(X)svb*^*108*^ and *Df(X)svb*^*108*^ x *svbBAC-dsRed*. A1 trichomes in the dashed boxes bounded by the two sensory cells, as indicated by the arrows, were counted. The bracket at the edge of the A1 stripe marks a region where trichome growth is exclusively covered by *DG3*, which disappeared with the deletion of *DG3* and did not recover with the introduction of *svbBAC-dsRed*. (D) The trichome number in larvae developed at 32 °C with *Df(X)svb*^*108*^ partially recovered to wild-type levels with the introduction of *svbBAC-dsRed*. The number of larvae counted was: 12 for wild-type *svb*, 28 for *Df(X)svb*^*108*^, 14 for *Df(X)svb*^*108*^ x *svbBAC-dsRed* and 13 for *Df(X)svb*^*108*^ x *DG3-lacZ*. One-sided Student’s *t*-test was applied for each individual comparison. In box plots, center line is mean, upper and lower limits are standard deviation and whiskers show 95% confidence intervals.

## Discussions

Transcriptional regulation is a complex and dynamic process which requires coordinated interactions between transcription factors and chromatin. Given the transient nature of these interactions, using multiple binding sites to ensure robust transcriptional regulation appears to be a preferred strategy among many developmental enhancers (Frankel, 2012; Perry et al., 2010). Genes such as *shavenbaby* add another layer of redundancy on top of this through long *cis*-regulatory regions containing multiple enhancers whose expression patterns overlap. Previous works have shown that this redundancy ensures proper development when systems are subjected to stress (Crocker et al., 2015a; Frankel et al., 2010; Osterwalder et al., 2018). However, the mechanism underlying this robustness was not clear.

In this work, we took advantage of the high-resolution imaging and analysis techniques we had developed to observe transcriptional microenvironments around transcription sites (Tsai et al., 2017) and investigated how the *DG3* enhancer confers robustness to the *svb* locus at the molecular level. Deletion of the *DG3* enhancer from *svb* did not lead to clear defects in ventral trichome formation unless the embryos were subjected to heat-induced stress, as shown here and previously with the deletion of “shadow enhancers” (Hong et al., 2008) for lateral *svb* expression (Frankel et al., 2010). Nevertheless, we observed that the *DG3*-deletion allele showed reduced transcriptional output even at normal temperature. Wild-type embryos did not show phenotypical defects under either normal or stressed conditions, but heat-induced stress still led to lower transcriptional output from the wildtype *svb* allele. Thus, having enhancers with overlapping expression patterns could ensure a sufficient margin to buffer against the negative effects of environmental stresses (Frankel et al., 2010; Perry et al., 2010). The transcriptional output of the wild-type *svb* is likely still sufficiently high when embryos are stressed, due to its high overall expression levels. However, the mutant *svb* locus, starting with lower transcriptional output even under ideal conditions, drops below a threshold and the system fails. As the system appears to tolerate significant drops in transcriptional output before phenotypical defects appear, direct observation of the intermediate steps in the gene expression process would be needed to dissect the mechanisms underlying transcriptional robustness. To achieve this end, future works would need to investigate how inactive genes interact with transcriptional microenvironments and also track microenvironment-gene interactions dynamically in living embryos in real time.

We previously observed that transcription sites of reporter genes driven by minimal *svb* enhancers tended to colocalize with the endogenous *svb* locus when it is transcriptionally active (Tsai et al., 2017). This is true also at the scale of entire *cis*-regulatory regions, as we observed that the *svb* locus does the same with *svbBAC*, which implies that they potentially share a common microenvironment. Previously, homologous regions have been shown to pair over long distances, both between homologous chromosomal arms(Lim et al., 2018), translocated domains and between different chromosomes (Gemkow et al., 1998). Our observations are in line with the suggested transcription-dependent associations of interchromosomal interactions (Branco and Pombo, 2006; Joyce et al., 2016; Lomvardas et al., 2006; Maass et al., 2018; Monahan et al., 2019). It is possible that such long range interactions are driven, or reinforced, through shared microenvironments.

We were able to partially rescue the *DG3*-deletion *svb* allele with *svbBAC*, which contains the *cis*-regulatory region of *svb* but not the *svb* gene itself. High-resolution imaging showed that colocalizing with a *svbBAC* increases the local Ubx concentration and transcriptional output of the *DG3*-deletion allele. This supports a mechanism where transcriptional microenvironments sequestered around large *cis*-regulatory regions can work in *trans* to increase transcriptional output of other genes, even on different chromosomes, so long as they are spatially close by and share similar transcription factor binding sites. Additionally, there may be a lower limit on the size of the regulatory region before it can sufficiently rescue a deficient microenvironment—the *DG3* minimal *svb* enhancer alone did not rescue deficiencies. It is possible such interactions require structural elements, such as insulator proteins (Lim et al., 2018) or other topologically associated elements (Furlong and Levine, 2018). This is consistent with recent findings for long-range interactions that are dependent on specific topologically associating domains (TADs) (Viets et al., 2018). Interestingly, the *svbBAC* could not completely rescue trichome formation at the tip of the ventral stripe in embryos with the *DG3*-deletion allele, where *DG3* alone provides coverage. It appears that microenvironment-sharing rescues through increasing the local concentration of transcription factors so that overlapping enhancers increase their expression to compensate, but the *cis*-regulatory region of a gene is still the determining factor if transcription can occur at all in specific cells.

We previously proposed that transcriptional microenvironments form across multiple enhancers through scaffolding interactions to ensure efficient transcription from developmental enhancers. By investigating the mechanisms of regulatory robustness using a *DG3*-deletion allele of *svb*, we have shown that transcriptional microenvironments could span multiple enhancers that share similar transcription factor binding sites. These microenvironments of transcription factors could form the core of transcriptional “hubs” that have been proposed to form through phase-separation (Cisse et al., 2013; Furlong and Levine, 2018; Ghavi-Helm et al., 2014; Mir et al., 2017). Thus, they add another layer of redundancy on top of multiple overlapping enhancers in a *cis*-regulatory region. This provides an extra margin of safety in transcriptional output, preserving phenotypical development even when environmental conditions are not ideal. Integrating multiple noisy and weak elements into a coherent and synergistic network would also reduce the variance stemming from the transient and stochastic transcription factor binding dynamics observed in eukaryotic cells (Cisse et al., 2013; Ghavi-Helm et al., 2014; Mir et al., 2017; Tsai et al., 2017). In sum, specialized transcriptional microenvironments could be a critical element to ensure that gene expression occurs specifically and consistently in every embryo. Given that shadow enhancers are widespread features of gene regulatory networks (Cannavò et al., 2016; Osterwalder et al., 2018), it is likely that high local concentrations of transcription factors are a widespread feature that provides an effective regulatory buffer to prevent deleterious phenotypic consequences to genetic and environmental perturbations.

## Materials and methods

### Fly Strains

All fly strains used have been previously described: *DG3-lacZ* (Tsai et al., 2017); *ubx1* (Crocker et al., 2015a); *HS::ubx-1*: (Crocker et al., 2015a); *Df(X)svb*^*108*^ (Frankel et al., 2010); *svbBAC-dsRed* (Preger-Ben Noon et al., 2018a); *diBAC-gfp* is CH322-35A16 EGFP tagged in VK37, covering D (Venken et al., 2009). Unless otherwise noted, they are generated from w^1118^ stock, which is referred to as wild-type.

### Preparing Drosophila embryos for staining and cuticle preps

*D. melanogaster* strains were maintained under standard laboratory conditions, reared at 25°C, unless otherwise specified. For heat-shock experiments, these conditions were followed: for staining with fluorescent antibodies, flies were allowed to lay eggs on apple-juice agar plates for 5h at 25 °C and then kept in an incubator at 32 °C for 7 hours before fixation; for cuticle preps, dechorionated embryos were kept at 32 °C until they emerged as larvae. *Df(X)svb*^*108*^ embryos/larvae with *svbBAC-dsRed* are readily discernable by the loss of *svb* and trichomes in the T1 segment (see Figure 2B, C).

### Cuticle preparation and trichome counting

Larvae collected for cuticle preparations were mounted according to a published protocol (Stern and Sucena, 2011). A phase-contrast microscope was used to image the slides. Ventral trichomes in larval A1 or A2 segments were counted in Fiji/ImageJ by find using the find maximum function (Schindelin et al., 2012; Schneider et al., 2012).

### Immuno-fluorescence staining of transcription factors and in situ hybridization to mRNA

Standard protocols were used for embryo fixation and staining (Crocker et al., 2015a; Tsai et al., 2017). Secondary antibodies labeled with Alexa Fluor dyes (1:500, Invitrogen) were used to detect primary antibodies. *In situ* hybridizations were performed using DIG, FITC or biotin labeled, antisense RNA-probes against a reporter construct RNA (*lacZ, dsRed, gfp*) or the first intron and second exon (16kb) of *svb*. See Supplemental table 1 for primer sequences. DIG-labeled RNA products were detected with a DIG antibody: Thermofisher, 700772 (1:100 dilution), biotin-labeled RNA products with a biotin antibody: Thermofisher, PA1-26792 (1:100) and FITC-labeled RNA products with a FITC antibody: Thermofisher, A889 (1:100). Ubx protein was detected using Developmental Studies Hybridoma Bank, FP3.38-C antibody at 1:20 dilution, DsRed protein using MBL anti-RFP PM005 antibody at 1:100, LacZ protein using Promega anti-ß-Gal antibody at 1:250 and GFP protein using Aves Labs chicken anti-GFP at 1:300.

### Imaging fixed embryos

Mounting of fixed *Drosophila* embryos was done in ProLong Gold+DAPI mounting media (Molecular Probes, Eugene, OR). Fixed embryos were imaged on a Zeiss LSM 880 confocal microscope with FastAiryscan (Carl Zeiss Microscopy, Jena, Germany). Excitation lasers with wavelengths of 405, 488, 561 and 633 nm were used as appropriate for the specific fluorescent dyes. Unless otherwise stated, all images were processed with Fiji/ImageJ (Schindelin et al., 2012; Schneider et al., 2012) and Matlab (MathWorks, Natick, MA, USA).

### Analysis of microenvironment and *svb* transcription intensity

Inside nuclei with svb transcription sites, the center of the transcription site was identified using the find maximum function of Fiji/ImageJ. A 40×40 pixel square region of interest (ROI) centered on the transcription site is then created. The integrated fluorescent intensity inside the ROI from the Ubx IF channel and the RNA FISH channel are then reported as the local Ubx concentration and the transcriptional output, respectively. The intensity presented in the figures is the per-pixel average intensity with the maximum readout of the sensor normalized to 255.

### Analysis of distances between transcription ‘spots’

Inside nuclei with svb and dsRed/GFP transcription sites, the centers of the transcription site were identified using the find maximum function of Fiji/ImageJ. The distance between the transcription sites were then computed using the coordinates of the transcription sites.

### Ubx ChIP profile

The ChIP profile for Ubx around the svb cis-regulatory region is from Choo et al. (Choo et al., 2011), using whole *Drosophila melanogaster* embryos between stages 10 and 12.

## Acknowledgements

The fly line containing *diBAC-gfp* (CH322-35A16 EGFP) was a gift from Schulze, Karen Lynn & Bellen, Hugo J. The *Df(X)svb*^*108*^ flies were a gift from Stern DL and the *svb*BAC flies were a gift from Preger Ben-Noon E and Frankel N (Preger-Ben Noon et al., 2018b). We thank Rafael Galupa and Nicolas Frankel for suggestions and discussions. We thank the entire Crocker lab for discussion and feedback. Albert Tsai is a Damon Runyon Fellow of the Damon Runyon Cancer Research Foundation (DRG 2220–15). Mariana R P Alves and Justin Crocker are supported by the European Molecular Biological Laboratory (EMBL).

## Figures

**Figure 1-figure supplement 1.**
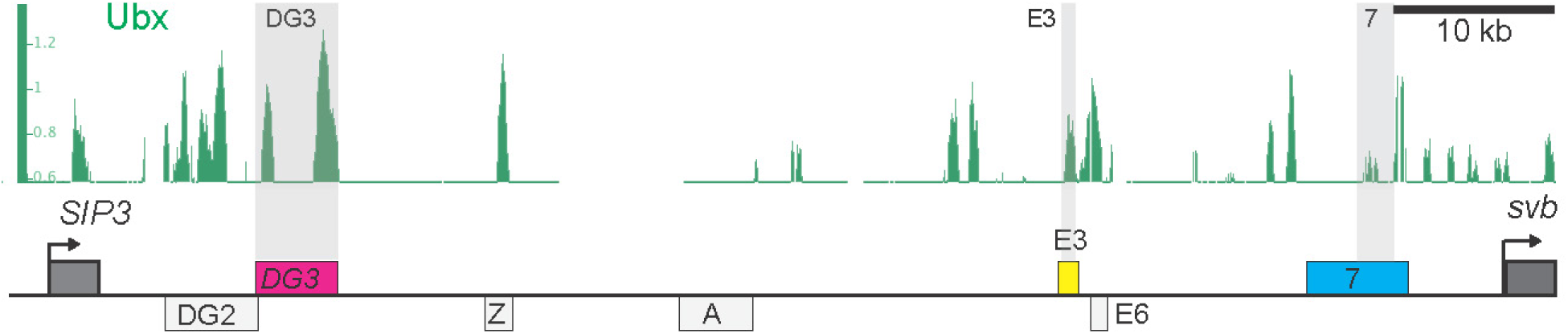
Ubx enrichment around the 3 ventral *svb* enhancers. ChIP experiment of whole embryos between stages 10 and 12 targeting Ubx shows different enrichment profile around the three ventral *svb* enhancers: *DG3, E3* and *7*.

**Figure 2-figure supplement 1.**
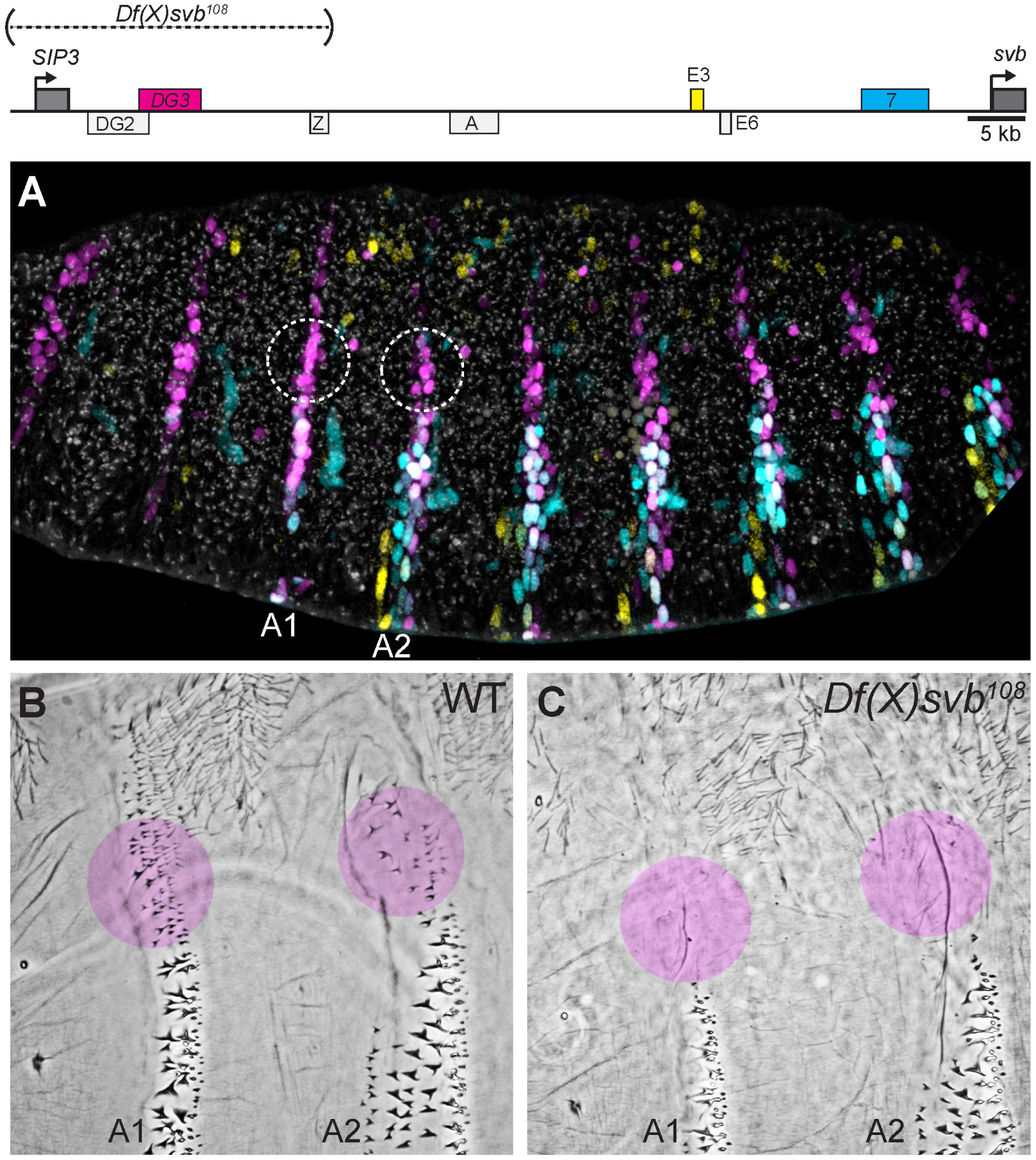
Loss of trichomes in regions exclusively covered by *DG3*. (A) Within the overall expression pattern of *svb, DG3* provides exclusive coverage in the circled regions in segments A1 and A2. (B & C) Even at 25 °C, where the overall trichome numbers in A1 and A2 for the *Df(X)svb*^*108*^ deletion mutant is indistinguishable from wild-type *svb*, the trichomes at the edge of the ventral stripe for A1 and A2 were lost. These patches correspond to the circled regions in panel A.

**Figure 4-figure supplement 1.**
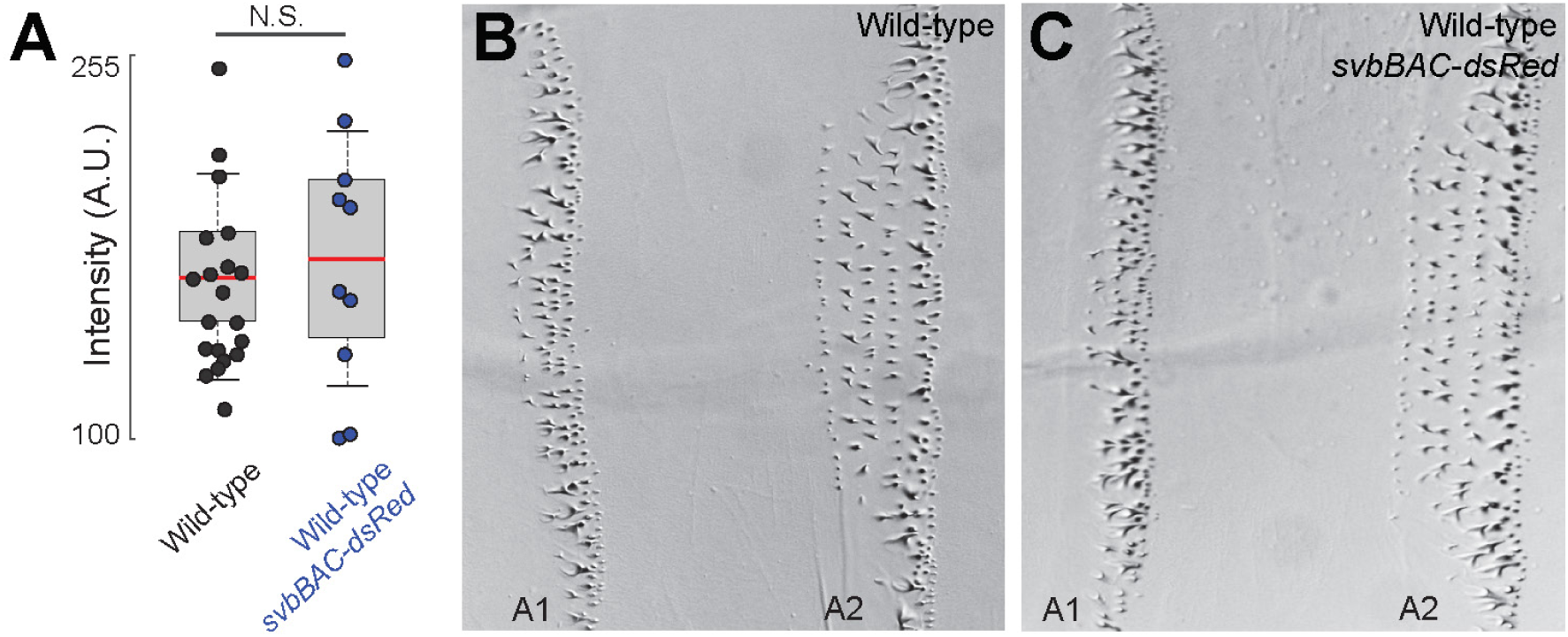
Introduction of *svbBAC-dsRed* to wild-type (w^1118^) does not change Ubx microenvironment and phenotype. (A) Ubx concentrations around *svb* transcription sties in wild-type (w^1118^) x *svbBAC-dsRed* embryos did not change compared to wild-type. The number of *svb* transcription sites quantified was: 20 for wild-type and 10 for wild-type x *svbBAC-dsRed*. One-sided Student’s *t*-test was applied for each individual comparison. We analyzed 3 embryos for each genotype. In box plots, center line is mean, upper and lower limits are standard deviation and whiskers show 95% confidence intervals. (B & C) The trichome phenotype along A1 and A2 did not change with the introduction of *svbBAC-dsRed* to wild-type (w^1118^).

**Supplementary table 1.**
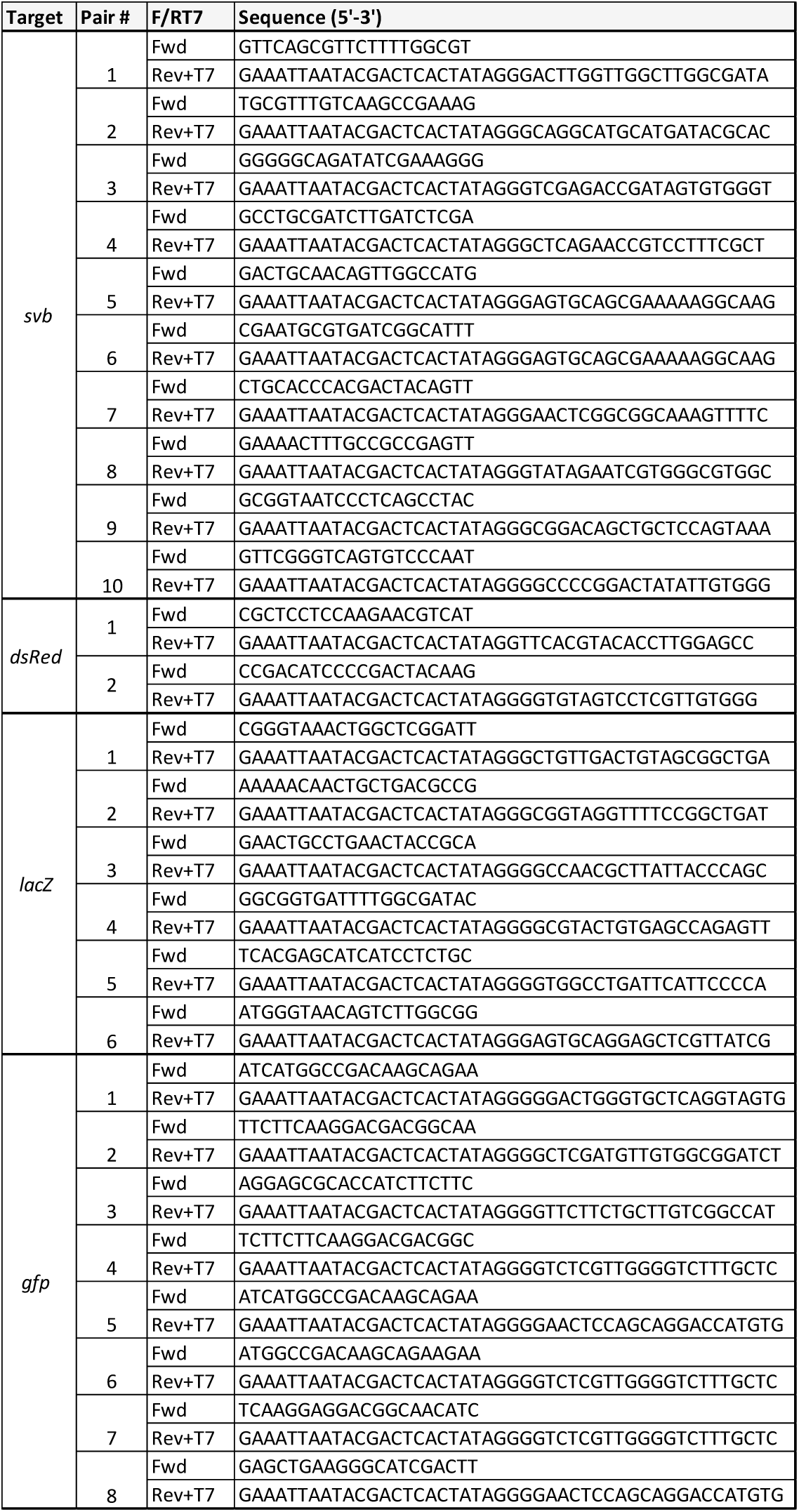
Primers for RNA-probe generation Sequences of primers for amplification of DNA to be used for generation of antisense RNA-probes. The targets - reporter construct RNA (*lacZ, dsRed, gfp*) and first intron and second exon (16kb) of *svb* - are indicated in the left column. Sequences are indicated for forward or reverse primers of each pair. Reverse primers include a T7 sequence for transcription with T7 RNA polymerase.

